# Patterns of molecular conservation along tooth development are only partly shaped by evolutionary pressures on tooth

**DOI:** 10.64898/2026.06.19.733320

**Authors:** Jérémy Ganofsky, Mathilde Estevez-Villar, Marion Mouginot, Sébastien Moretti, Marion Nyamari, Marc Robinson-Rechavi, Sophie Pantalacci, Marie Sémon

## Abstract

Although it is well established that certain stages of development are molecularly more conserved than others, the reasons for this phenomenon remain largely unknown. We study molecular conservation in the development of an organ, the molar, by comparing the temporal profiles of expression in mice and hamsters. We find that the cause of conservation of expression and of coding sequences changes over molar development. Gene expression levels display a classical increase of divergence as development progresses. In terms of genes expressed, the composition of early and late stages is better conserved and enriched in pleiotropic genes, yet each stage mobilizes different sets of pleiotropic genes, cell division for bud growth and secretion for tooth mineralization. Moreover similar patterns of higher divergence of gene sets and of coding sequences at mid development, are caused by different biological phenomena, in that case heterochronies and blood colonisation respectively. In conclusion, the patterns of molecular conservation in developing molars are shaped by a combination of processes intrinsic to the teeth, and by negative and positive selection on functions which are mostly extrinsic to the teeth. This is likely translatable to explain molecular conservation patterns in many other biological systems.

**AUTHOR SUMMARY:** For species to evolve different adaptations to different life styles, their anatomy has to evolve correspondingly. This in turn implies evolution of the embryonic development of anatomical structures. Notably, tooth shape can evolve rapidly as an adaptation to different diets. Mice and hamsters are closely related rodents who yet differ in the shape of their molars, and thus in their development. In this study, we investigated why the genes active in molar development are more or less similar between the two species from early tooth bud to fully formed embryo molar. We found that early and late molar development were slow evolving, while mid-development was evolving faster. But surprisingly, this was in part due not to tooth evolution, but to the involvement of genes which are active in other processes in the body. For example an influx of immune cells also brings fast evolving immune genes. This helps us understand better the complexity of causes of apparently simple evolutionary patterns.

## INTRODUCTION

Transcriptomics, which allows genome expression to be sampled and compared at different developmental stages between species, has become an effective quantitative approach to measure embryonic molecular conservation. For over a decade, transcriptome studies have been measuring molecular conservation during embryogenesis and organogenesis, and then interpreting the results in terms of adaptation and environmental or developmental constraints (eg. [1–9]). Part of the research field is moving toward single cells and cell lineage dynamics [10–12].

An implicit assumption of this approach is that the conservation patterns of genes expressed in specific organs or developmental stages mostly reflect the evolutionary processes affecting these organs and stages. Yet many genes are pleiotropic, because they are widely expressed for house-keeping functions or they have specific functions which are expressed in a diversity of organs, or they have several different organ-specific functions. In all of these cases, genes might be strongly impacted in their evolution due to their activity in other conditions than the organs or stages directly under study.

Transcriptome studies of evolution of development have reported different patterns, using modular *versus* global transcriptome approaches, as well as different indexes and sampling strategies. For global patterns, following a first landmark study in drosophila several analyses have directly measured divergence in temporal expression profiles [6]. Another pioneering study introduced the transcriptome age index (TAI) that combines phylostratigraphy and stage-specific gene expression to reflect the evolutionary age of the transcriptome [13]. An early study of modules of expression in development also used this TAI, as well as other metrics [14].

Three main patterns of conservation have been observed. The first, corresponding to the “developmental hourglass” model, suggests that divergence is greater in early stages, then falls to a waist of conservation at mid-development and then increases once more throughout the remaining stages e.g. [6,11,12,15–19]. More recently, an hourglass pattern was discovered by comparing mouse and zebrafish development at single-cell level, but with an offset in the minimum of conservation, placed at the neurula stage by the transcriptome divergence index and at the pharyngula stage by the TAI index [10]. The second pattern, corresponding to the “developmental funnel” model, suggests divergence gradually grows from the early stages, that are most conserved, across the remaining developmental stages. This has mostly been observed when sampling starts later, and thus misses the early part of the hourglass [20,21]. The third pattern, the “inverse developmental hourglass”, is less frequently seen. It features conservation at early and late stages of development and a rise in divergence at mid-development [22–26]. Interestingly, for the same dataset, different indexes may give different results, such as a funnel of gene age or of duplicability, and an hourglass of gene regulation or of sequence conservation [14,27].

Beyond describing patterns, much work has been devoted to deciphering the evolutionary processes that create them, and their relation to environmental adaptations or internal causes, such as development or genetic properties. In the “funnel model”, it was proposed that very early and late stages are more subject to positive selection linked to ecological adaptations [28]. In the “developmental hourglass model”, it has been argued that embryonic development is particularly conserved at the phylotypic period, when gene regulatory complexity is higher and highly conserved key developmental genes orchestrate body plan organization [29–31], while early and late development are more permissive to the fixation of mutations. The importance of internal genetic constraints such as pleiotropy has also long been emphasized [32–34]. In most functional genomics studies, pleiotropy is operationally defined for a gene by compiling the different organs, developmental stages, or cell types in which it is expressed [21,35]. Thus defined pleiotropic genes are highly mobilized during early development, which is thought to reduce the number of mutations available for positive selection. The effects of functional constraints and adaptive changes are nonexclusive [36–38]. Accounting for pleiotropy is important not only as a global metric leading to constraints, but also because the development of a given organ can mobilise gene modules whose evolution is mostly impacted by their function in another organ, potentially outside of a developmental context. Working at the level of the whole embryo in distant species makes it difficult to disentangle all these effects. For this we need closely related species, organs with well known evolution and development, and to combine whole transcriptome approaches with modular analyses.

Here we chose the upper and lower molars, in two rodent species, the golden hamster (*Mesocricetus auratus*) and the mouse (*M. musculus*), that diverged ≈26 million years ago [39]. Molar tooth development is well described and mechanistically well understood [40–43]. The upper molar has acquired a new shape in the mouse, with two supplementary cusps. We have previously shown this arose from changes in cell proportions in early development, resulting in formation of supplementary cusps in late development [44,45]. The lower molar shares some of these developmental changes, but only the upper molar develops the additional cusps.

We have extended a previously published transcriptome time series, and established patterns of conservation of expression, adaptive and purifying selection in the coding sequence and gene age throughout molar development. We find that different patterns of molecular conservation characterize the evolution of tooth development due to the influence of pleiotropy, heterochrony, and old patterns of homology on different levels of organisation, from individual genes to global expression patterns.

## RESULTS

### Transcriptome profiling of mouse and hamster first molar morphogenesis

To compare the temporal dynamics of gene expression between mouse and hamster molar development, we gathered 103 bulk RNA-seq samples from a time series of growing upper and lower molar tooth germs (36 samples were collected specifically for this study to complete our previously published data, S1 Table, all taken from female embryos). We selected tooth germs from Embryonic day (E)12.5 to PostNatal day (PN)2 mice because this is the period during which the major events of morphogenesis occur, from bud, to cap, to bell stage until differentiation and enamel/dentin secretion. For the hamster, we selected the stages of dental development over a period as wide as in the mouse (from E11.0 to PN2). In order to compare stages between species, it is necessary to closely observe the entire sequence of morphogenesis because mice and hamsters develop at different rates, both in terms of the length of the pregnancy and in terms of the overall advancement of development. We did this by leveraging our earlier comparative research of tooth and palate development in these two species [46–49]. We used the fact that embryo weight is well correlated with its developmental age (S1 Text, Fig 1A, S1 Fig) and we observed the presence and number of tooth and cusp signalling centers in the isolated epithelial part of the molar germ, so that the progression of morphogenesis can be fully appreciated.

**Fig 1.**
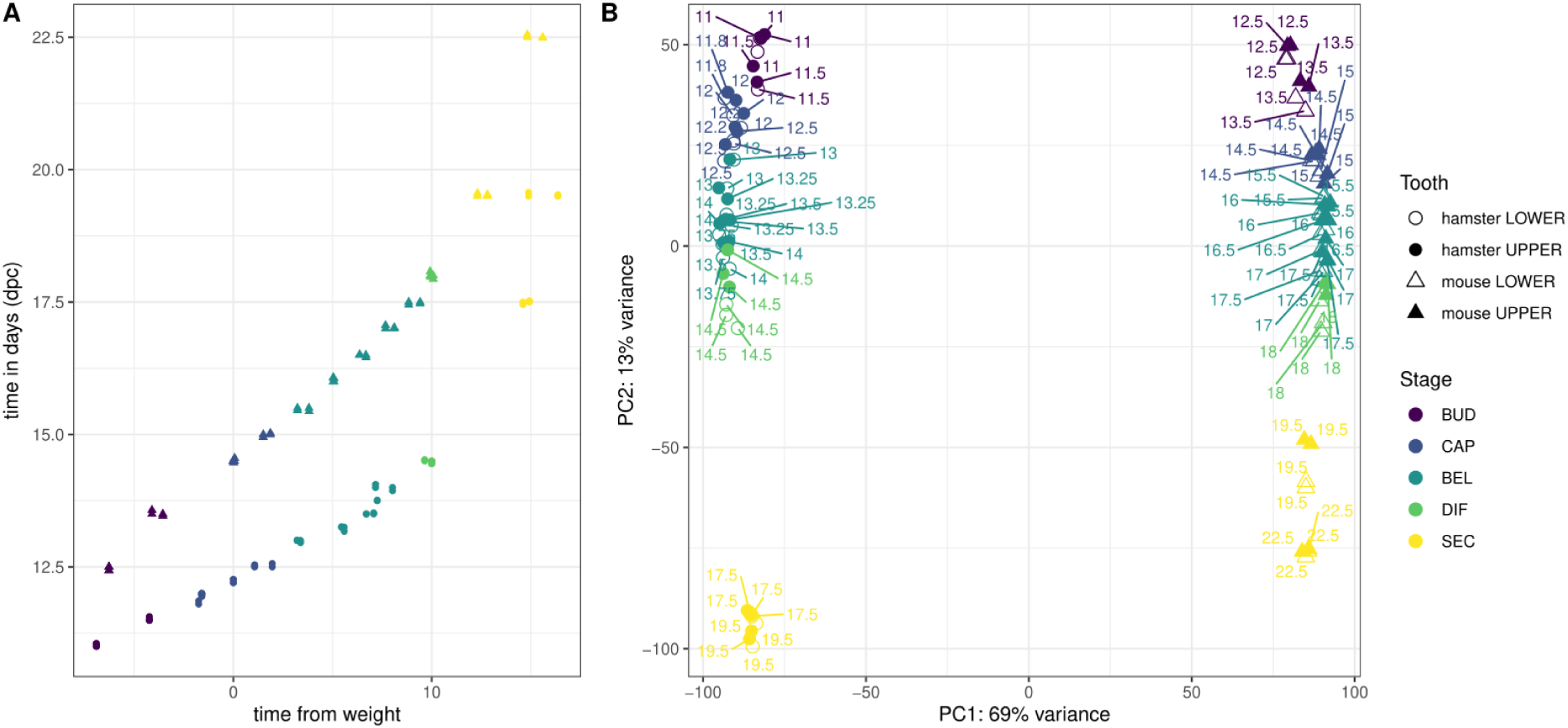
Transcriptome profiling of molar development. (A) Mouse and hamster samples (triangles and circles) in days post coitum (dpc) and in relative time estimated from embryonic weight. (B) PCA based on normalized log-transformed TPM expression levels, for 11,342 1:1 orthologs across mouse and hamster. Samples are colored per morphological stage, shapes correspond to species and upper and lower molar samples.

To obtain the patterns of evolutionary conservation we first needed to group samples according to their developmental stages. We attributed samples to five acknowledged morphological stages characterizing tooth development: bud, cap, bell stage, differentiation and secretion (Fig 1, S1 Table, S1 Text, S2 Fig). During the *bud stage (BUD)*, the epithelium invaginates while recruiting mesenchymal cells. During the *cap stage (CAP)*, the epithelium progressively wraps around the condensed mesenchyme, which delineates the future crown. During the *bell stage (BEL)*, the cusps tips are patterned, which initiates the ameloblast and odontoblast specification process. During the *differentiation stage (DIF)*, these cell types continue to differentiate, with cusp tips and oldest cusps displaying the most advanced cell types, while other tooth regions remain less advanced. Ultimately, during the *secretion stage (SEC)*, these cells mature and secrete a matrix that mineralizes as either enamel or dentin. The relative times of these morphological stages are coherent between species (Fig 1A, S3 Fig shaded bands). Of note, the stage where the offset is maximal is the secretion stage, because to be conservative, we selected day 0 and day 2 post-birth for both species despite an expected offset, since physiological changes at birth may have systemic effect and impact tooth transcriptome (see S3 Fig and S1 Text).

### Shifts in expression peaks are less frequent for early or late peaking genes, while divergence in expression levels is greater in late stages

To study the evolution of organogenesis, we believe it is important to study temporal expression profiles and not just global expression levels. Indeed, over development, there is a mix of genes which are specific to some developmental stages, and of genes which are not (sometimes called “housekeeping genes”, which are expressed generally without having a stage-specific functional role). We think, following [50], that even though genes may potentially be used outside of the stages in which they are most expressed, the shift in expression level is a good way to determine how involved a gene is in various stages. Besides, genes that show variation in expression levels during the course of development, may play an important role even if they have low expression levels. Grouping genes in temporal modules therefore allows us to identify and study the evolution of processes which characterise specific stages.

The simplest way to build such modules would be to group genes specifically expressed in a single stage, but this represents very few genes (S2 Table). This reflects the biology of molar formation, where the same morphogenetic mechanisms are iterated for cap and sequential cusp formation (genes shared during BUD, CAP and BEL stages), and the same cell specification/differentiation processes occur at very different times in different parts of the tooth (genes shared during BEL, DIF and SEC stage, with differentiation of cells deep in the valleys being strongly delayed compared to the cusp tip, see more explanation in S1 Text). Simple presence/absence calls are therefore not applicable to our data to study evolutionary conservation.

We decided instead to classify genes according to quantitative variation in expression during development. Applying a flexible clustering approach with semi-supervised starting conditions [51,52] to each species and tooth type, we clustered together genes whose expression peaked at the same stage. We call these “clusters of coexpression” further on. We associated about three-quarters of the genes with a cluster, confirming the labile nature of gene expression during development, even in a relatively short period of morphogenesis and in a single organ. The distribution of cluster sizes is similar for all teeth, with more genes associated with early and late stages than with intermediate stages, again in consistency with the biology of molar development (Fig 2A).

**Fig 2.**
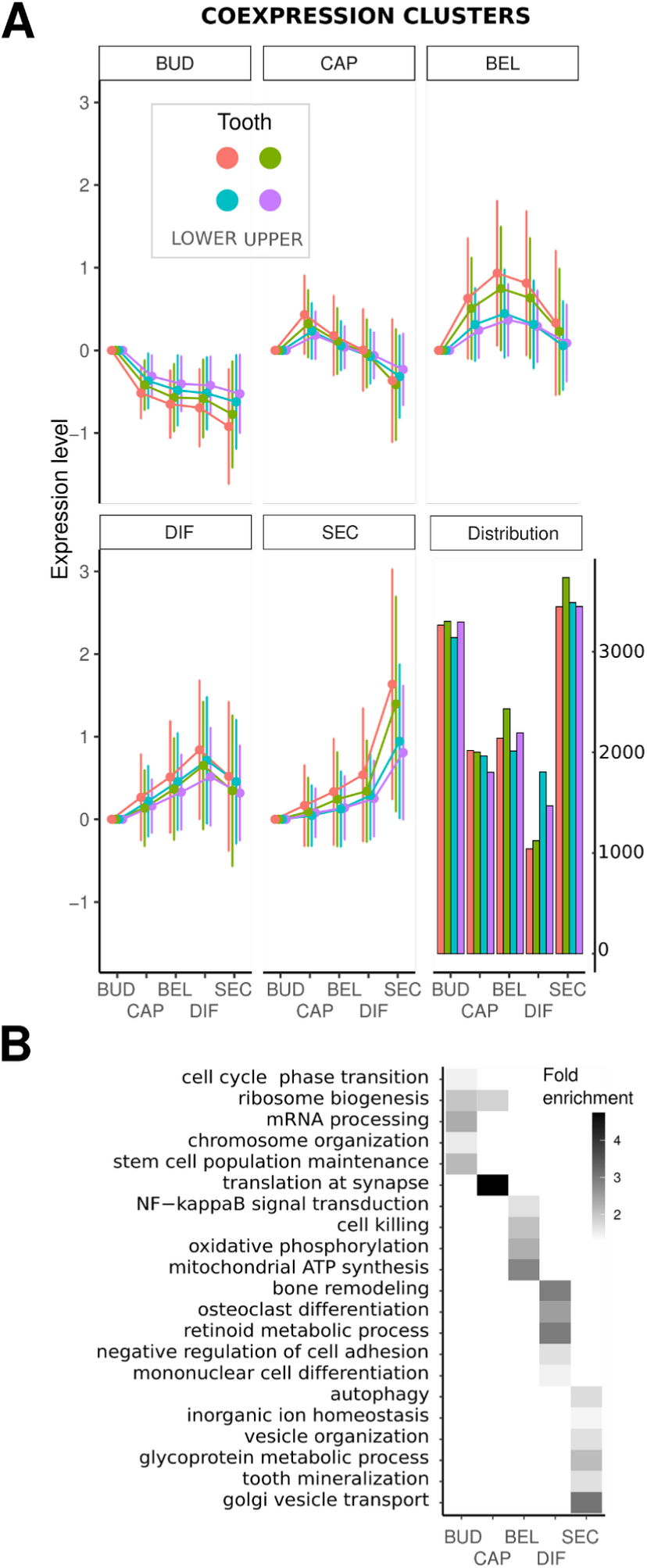
Clusters of co-expressed genes with an expression peak at a specific stage of dental development. Coexpression clusters defined by soft clustering of expression values of each species and tooth type. Each cluster corresponds to a set of genes with an expression peak at a specific stage of dental development. (A) The average profiles of each cluster are represented, with expression levels centered at the value of the first stage and error bars showing standard deviations. Clusters are ordered according to their temporal profiles. Distribution of cluster sizes is given. Genes not associated with a cluster are not represented. (B) Summary of the GO enrichment analysis for the mouse lower molar coexpression clusters; enriched GO terms were categorized as shown in S4 Fig, a maximum of 5 representative terms are shown with their fold enrichments.

To associate these clusters with biological functions, we performed a gene ontology (GO) enrichment analysis on each cluster (Fig 2B, S4 Fig, S5 Fig). In the mouse lower molar, we observed an initial phase associated with cell cycle, protein synthesis, and metabolism (BUD), followed by an intermediate phase with similar (yet more modest) enrichments plus indicators of nerve colonization (CAP), then by a phase associated with cell-cell signaling and mitochondrial respiration (BEL), a phase associated with mineralized tissue formation and blood cells (DIFF), and finally a phase associated with autophagy, secretion, and mineralisation (SEC). This temporal sequence is in agreement with the succeeding processes that contribute to molar formation, as a mineralized but also richly vascularized and innervated organ [53]. Similar enrichment terms of GO terms were observed for the other molars.

Having clustered together genes whose expression peaked at the same stage in the time series of each tooth, we systematically compared the gene content of these clusters to determine whether the same genes peak at the same stage in each tooth. We measured their overlap in terms of enrichment (fold change) relative to random expectation (see Methods). The largest overlaps were observed between upper and lower molars of the same species (S6 Fig), where clusters corresponding to the same stage have 60-70% of genes in common, which is 3 times more than would be obtained by chance. This is consistent with previous results on lower and upper molar transcriptome coevolution [54].

Although overlap was also observed between species, there were marked differences in gene content (Fig 3), again consistent with the rapid change in developmental expression observed in a previous study [55]. A third of the genes associated with a cluster whose expression peaks at a given stage in the upper molar of the mouse was associated with the cluster peaking at the same stage in the upper molar of the hamster, which is 50% more than expected (Fig 3C) but still a minority of the genes. The largest overlaps are observed for clusters with expression peaks at early and late stages. The difference is statistically significant: its magnitude far exceeds the bootstrap intervals (segments on the solid lines) and it exceeds the temporal variation observed on the shuffled data (dashed lines, Fig 3C,D).

**Fig 3.**
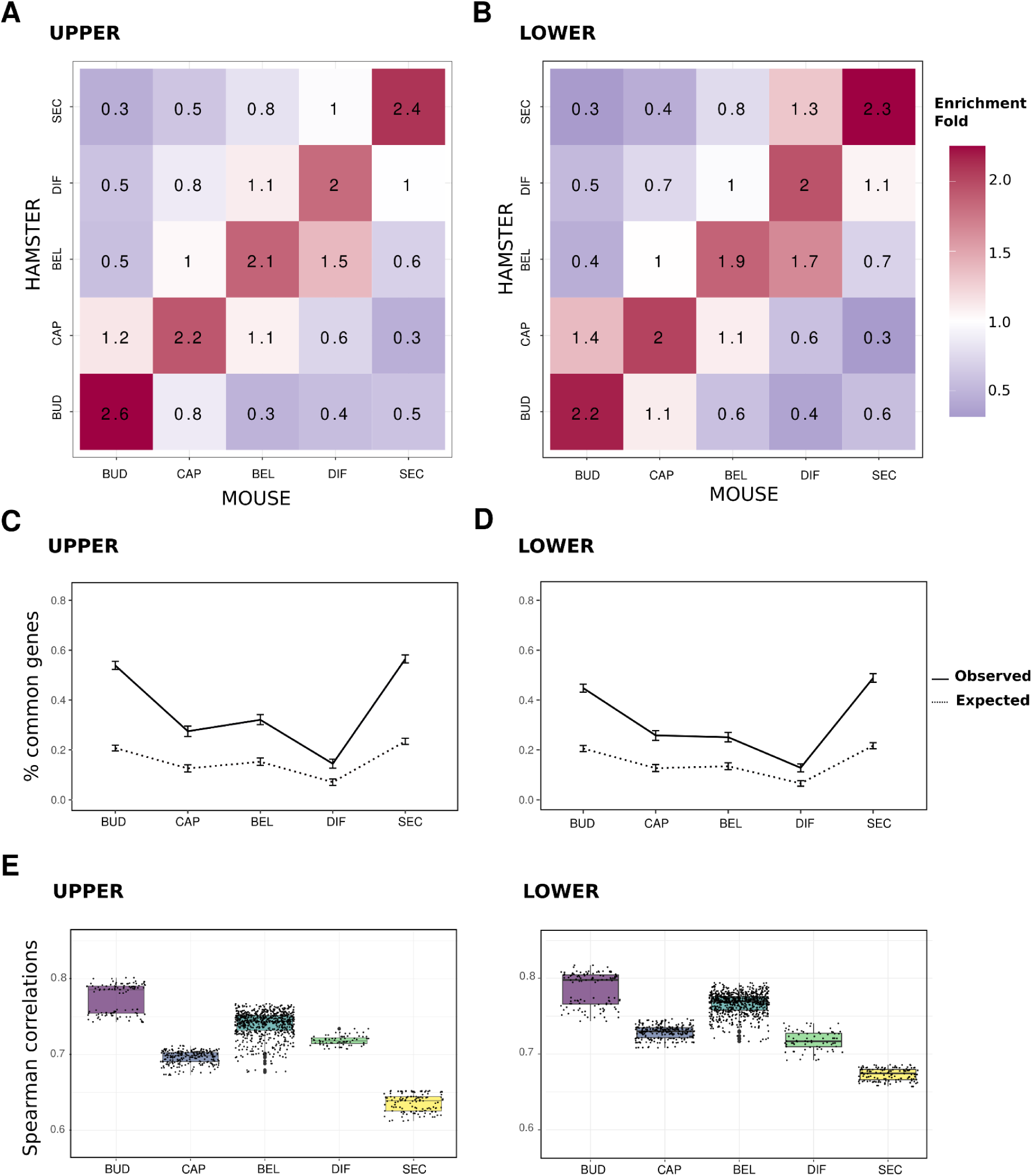
Conservation of expression gene clusters between species. Each cluster corresponds to a set of genes with an expression peak at a specific stage of development in a given tooth and species. Number of genes shared between hamster and mouse coexpression clusters, expressed as a fold enrichment relative to equally sized random clusters, for the upper (A) and the lower (B) molars. Clusters are ordered according to their temporal profiles. Proportion of genes shared between hamster and mouse coexpression clusters peaking at the same stage in both species, in the upper (C) and lower (D) molars. Intervals show the 95 percentiles obtained by 1000 bootstrap resamplings. Expected values obtained by randomly shuffling genes between clusters (dashed lines). (E) Expression conservation at each stage for the gene clusters peaking at this stage as measured by Spearman correlations between pairs of samples in mouse and hamster.

Because genes may peak at the same stage in mouse and hamster but still have diverged in terms of expression levels, we computed distances based on Spearman correlations (Fig 3E). We observed that expression levels were conserved for the BUD stage, divergent for the SEC stage, and intermediate in between.

The previous approach relies on gene clustering. To study divergence with all the genes and a continuous manner, we modelled temporal profiles by using spline models as in [44,45]. We observed that the timing of the peaks of expression were most conserved in the early and late stages, which was consistent with the conservation of peaks in the clustering approach (S7 Fig, S8 Fig). We observed that the pattern for the divergence in the levels of expression was also consistent with the clustering approach (S9 Fig). Therefore with this independent method, we recovered similar patterns as with stage-specific gene clusters.

Altogether, we observed the stages of morphogenesis have diverged differentially in terms of gene content and in terms of gene expression level of these gene sets. While the BUD stage shows high conservation with both indexes, the SEC stage mobilized the same gene content but these genes have diverged in expression levels. Other stages show medium behavior, also more variable between teeth and between approaches.

### Constraints on coding sequences are higher for both genes expressed at early and at late stages

To assess potential differences in functional constraints during development, we compared the selective pressure operating on gene coding sequences between coexpressed gene clusters. We quantified the mode and strength of selection by using dN/dS ratios.

We obtained substitution rates per gene, and averaged them per coexpressed gene cluster for each tooth, to compare average values of dN/dS ratios between clusters associated with different stages. These dN/dS values are low overall, showing generally purifying selection on genes expressed in tooth development; they are typical of the range of the whole genome (S10 Fig, median: 0.1303, mean: 0.173). Yet there is significant variation between the genes expressed at different stages of molar development, which sheds light on the variation in selection on these genes. In particular, the ratios are significantly lower at early and late stages than at intermediate stages. This is true for all tooth types, with a maximum ratio (i.e. weakest constraint) for the differentiation stage in the mouse, and for the bell stage in the hamster. This creates an inverse hourglass pattern of purifying selection levels operating on coding sequences (Fig 4). Although gene modules are quite large (Fig 2), they do not encompass the whole transcriptome, and one may also consider that genes with an expression peak at a given stage do not alone determine the evolutionary conservation of that stage. To control for this, we computed a transcriptome index of purifying selection on coding sequence, calculated as the dN/dS of each gene weighted by its expression at each morphological stage (following [56]). This gave qualitatively similar patterns, with stronger signs of purifying selection at BUD and SEC stages than in the middle organogenesis (S11 Fig and S12 Fig).

**Fig 4.**
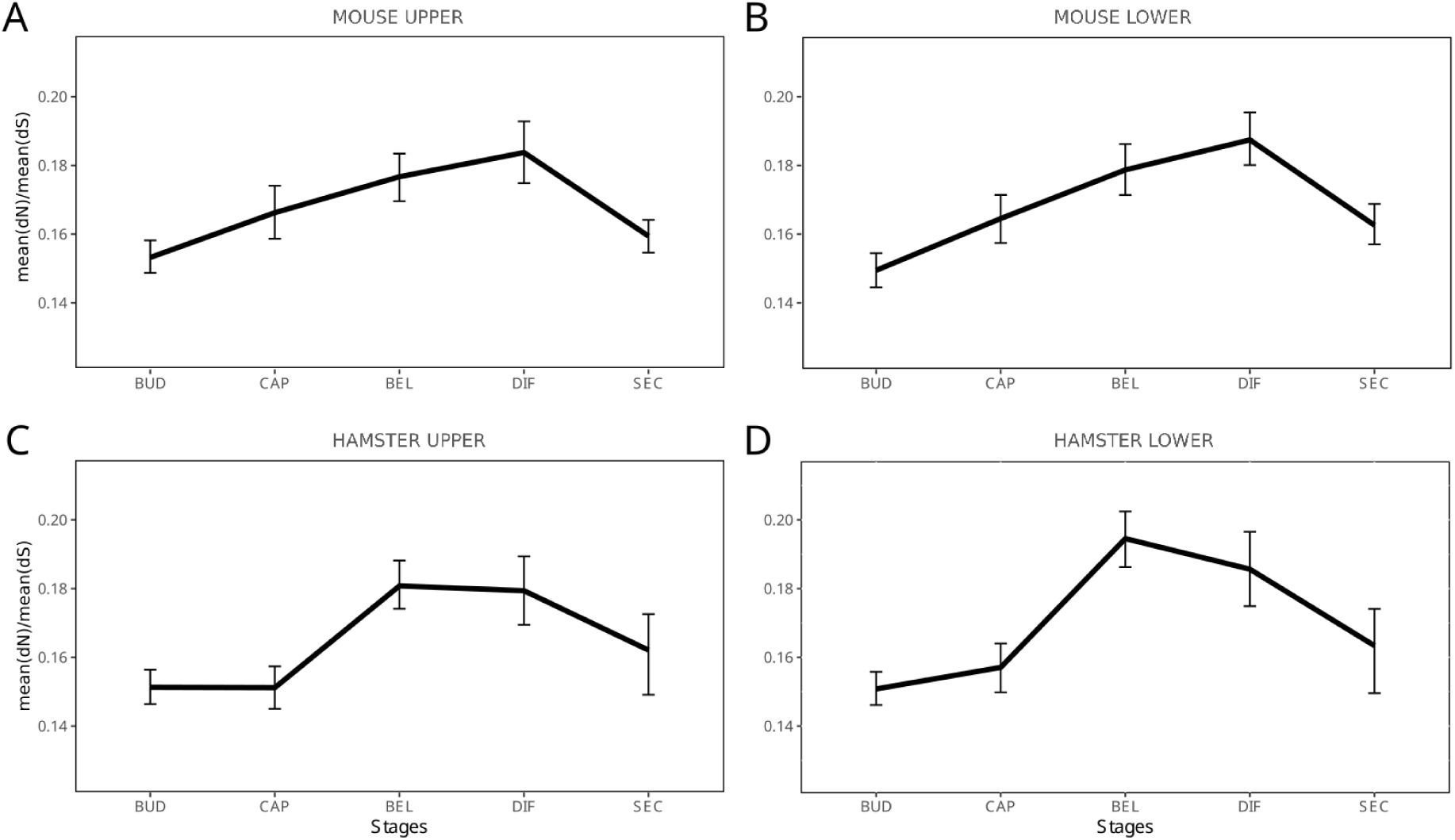
Selective pressure acting on gene coding sequences for each coexpressed gene cluster. Average pairwise dN/dS between mouse and hamster were computed as mean(dN)/mean(dS). Observed values are represented by a black line, intervals show the 95 percentiles obtained by 1000 bootstrap resamplings.

We verified this pattern both for genes with conserved and with non-conserved expression peaks between pairs of molars (S13 Fig). In addition, genes with a conserved expression peak have significantly lower dN/dS values in most cases (S3 Table), indicating that conserved expression profiles are correlated with stronger purifying selection on their coding sequence. One possibility is that dN/dS profiles reflect different levels of constraints on tooth development. Another possibility is that this reflects the temporal usage of different sets of genes, whose dN/dS profiles are dictated by their role in many parts of the body, i.e. by their pleiotropy (see below).

### Adaptation in coding sequences tend to increase in mid molar development genes

Next, we evaluated the relationship between positive selection and developmental stages by examining the genes identified as carrying signatures of positive selection. We computed the proportion of genes with significant evidence of positive selection on several branches of the rodent tree (q-value < 0.2, S4 Table). We found 18 genes on the *Muridae* branch (out of 11460 tested genes), 60 on the *Murinae* branch (12919 genes), 49 on the *Mus* branch (13257 genes), and 36 on the *Cricetinae* branch (10936 genes). Except for one gene, Csf1r, these genes have no clear role in tooth development [57], but many are involved in immunity (S14 Fig, S15 Fig). This suggests that they were selected for other purposes than for tooth function, even though we cannot exclude that they may later serve immunity in the functional adult tooth. We grouped the genes by their associated stage and looked for pathway enrichments: bell/differentiation genes displayed significantly more interactions than expected with an enrichment for inflammatory response (PPI enrichment p-value, 0.019, N=47) while the genes associated with other stages did not (for bud/cap, N=42 genes and p-value=0.163, for secretion, N=30 and p-value=0.134, S14B Fig).

To compare the magnitude of positive selection across different stages, we used the likelihood ratio (Δln*L)* of the models with and without positive selection, following [58,59]. A branch in a gene tree with a higher Δln*L* value indicates higher evidence for positive selection for this gene over this branch. We computed ΔlnL for different branches of the species tree, and used gene coexpression clusters relevant for each branch (see Methods). The amount of positive selection increases from the early stages and is maximum at the bel/differentiation stages. This trend is particularly marked on the *Mus* branch but visible in all branches (Fig. 5, S16 Fig for transcriptome index where each gene is weighted by its expression).

**Fig 5.**
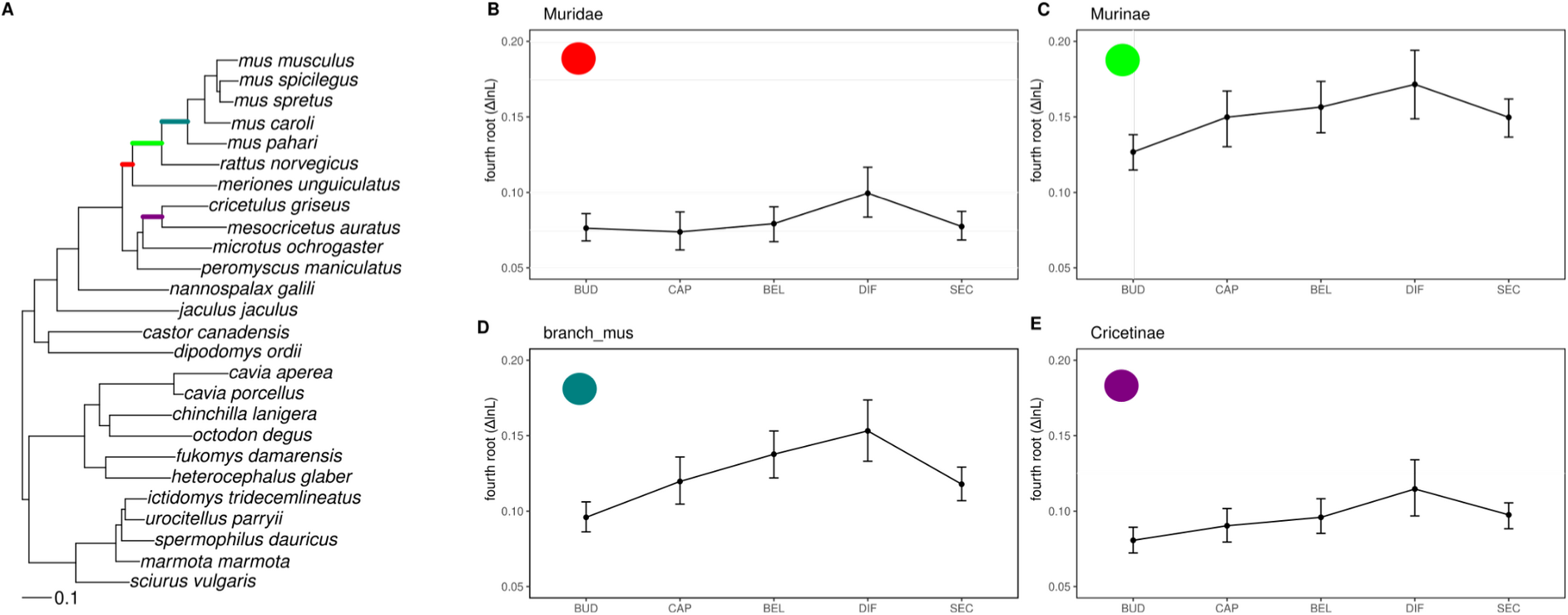
Positive selection for genes with expression peaks at different stages of development, in four branches of the rodent tree. Each plot shows the fourth root of the likelihood ratio (ΔlnL) of the models with and without positive selection, taken from Selectome database. Average for coexpressed gene clusters associated with each stage of development are represented. Confidence intervals obtained from 1000 bootstraps. Branches used to compute positive selection are indicated on the rodent tree in A, and correspond respectively to B: Muridae, C: Murinae, D: Mus, E: Cricetinae. Gene clusters correspond to mouse upper molar (B-D) or to hamster upper molar (E). Species tree in panel A pruned from Ensembl Compara.

It is well known that immune-related genes exhibit distinctive molecular evolution patterns and, more specifically, that they significantly contribute to the sequence-based signals of positive selection [60]. As a control, we therefore eliminated these genes from our analyses to examine how they affected the signals of evolutionary conservation. As anticipated, the positive selection signal is significantly reduced and almost completely absent (S16 Fig); however, the other patterns, whether they relate to expression levels or purifying selection on coding sequences, do not alter (S18 Fig, S19 Fig, S20 Fig, S21 Fig) with a stronger evolutionary conservation seen in mid-organogenesis.

### Variation in expression levels and pleiotropy explain the developmental pattern of sequence conservation

The influence of different parameters such as levels and breath of expression on the rates of protein coding sequence evolution has been extensively studied in many organisms [61–65]. In mouse, genes with high expression levels tend to evolve under strong purifying selection, although the effect size is small when confounding factors are taken into account [66]. We confirmed this negative correlation in our data, indicating that at all stages, expression in molar development correlates with the intensity of purifying selection on the coding sequence of genes (S22 Fig).

We next tested whether the relationship between dN/dS and expression level impacted the patterns we observed in Fig 4. We split genes in quantiles according to their maximum expression level and computed mean dN/dS separately for each expression category. We confirmed that dN/dS was higher for genes in low expression quantiles than for genes in high expression quantiles (S23 Fig). Within expression quantiles, there was a slight variation in dN/dS over the different developmental stages, but it was not significant. Hence the “inverse hourglass” pattern of conservation is seen at the global level but vanishes within subsets of genes with similar expression levels. The only remaining pattern is seen in the low expression quantile. Therefore, the global pattern must stem from the fact that the groups of genes expressed at different developmental stages have different properties, in particular in terms of expression levels.

The proportion of highly expressed genes indeed varied drastically between developmental stages (Fig. 6A-D). At the early stage (bud), the vast majority of genes belonged to the class with high or medium-high expression levels (70%), whose sequence evolves more slowly on average. In the intermediate stages, the proportion of highly expressed genes dropped (about 40%) and increased again at the last stage (about 60%). The proportion of low-expression genes followed of course a complementary pattern. Thus, the inverse hourglass pattern of sequence conservation is partly driven by the fact that the composition in genes with different levels of expression vary between different stages of development. Having found that expression level was a hidden factor explaining the sequence coding pattern led us to check if it could also explain the expression pattern. The proportion of conserved co-clustering genes is higher for highly expressed genes than low expressed genes and this is especially true for BUD and SEC stages (S24 Fig). Thus this follows, but does not create the pattern.

**Fig 6.**
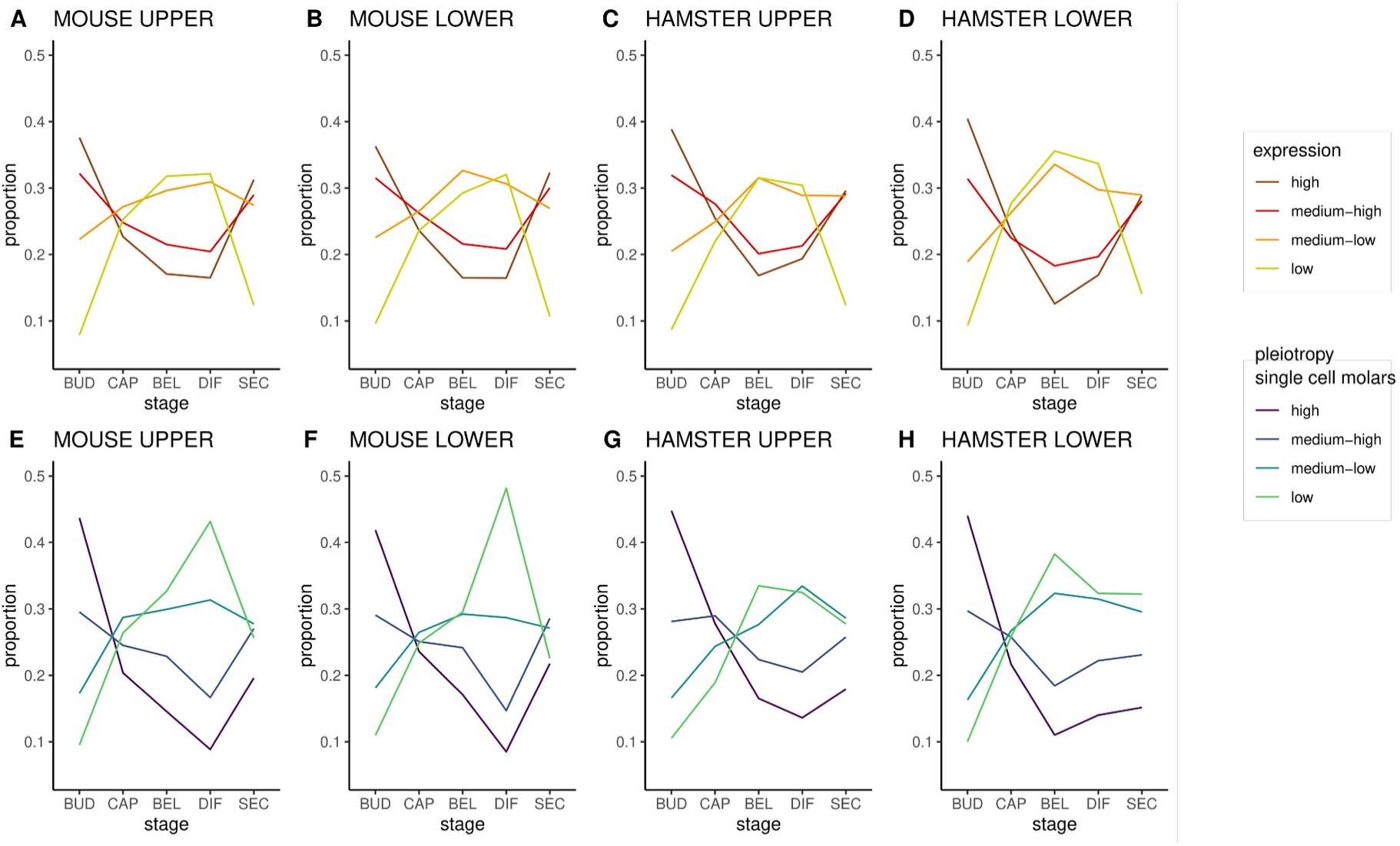
Genes expressed at different stages of development harbor different levels of expression and of cell expression specificity in molar. A-D : Genes were split in quantiles according to their average expression level. The proportion of genes associated with each quantile is drawn for each developmental stage and for each molar. Quantile boundaries are the same for all stages, and are computed separately for each molar, respectively quantile 1 to 4 in TPM for mus mx: 1.7, 20.9, 74.9,; for mus md: 1.6, 20.9, 74.8 for ham mx: 1.6, 17.2, 64.2; for ham md:1.5, 17.1, 63.9. E-H: Genes were split in quantiles according to their pleiotropy. The proportion of genes associated with each quantile is drawn for each developmental stage and for each molar. Quantile boundaries are the same for all stages, and were computed on tau values measured on mouse molar single cell transcriptome, with three threshold values ranging from low to high pleiotropy (25% 0.37, 50% 0.61, 75% 0.85).

A same level of expression in a whole organ may be due to very high expression in a limited subset of cells, or to more moderate expression, but in a very large number of cells [67]. To assess this, we measured for each gene an index of expression specificity, the Tau index, based on the different cell types present in a developing tooth germ. We used published single-cell RNA-seq (sc-RNAseq) data for embryonic mouse lower molars, which we clustered in 23 cell populations [68]. We split genes in quantiles according to their level of cell-specificity and compared their proportions across tooth development (Fig. 6E-H). At the bud stage, we observed that more than 40% of genes are pleiotropic (here: expressed in many cell types) and merely 10% are cell-specific. After that, the relative proportions reverse with about 10% pleiotropic genes at bell and differentiation stages. This mirrors the pattern observed for the quantiles of expression. In the latest stage however, the proportion of highly expressed genes increases again, while the proportion of highly pleiotropic genes remains modest. Hence, genes expressed in early and late stages tend to have different properties. Genes expressed in the early stage are expressed at a higher level in many cell types, while some genes expressed in the latest stage are also expressed at a high level but concentrated in fewer cell types (S25 Fig).

Because expression specificity was estimated from a single scRNA-seq sample, it may not represent all cell types present in all stages of the molar timeserie. Incisors grow continuously in mice, therefore presenting in a single organ many of the tooth cell types and for these, spanning stem cell differentiation gradients. We measured cell expression specificity on another scRNA-seq dataset from adult mouse incisors and obtained similar results as with the index calculated on molar scRNAseq data (Fig 7, 17 major cell subpopulations, [69]).

**Fig 7.**
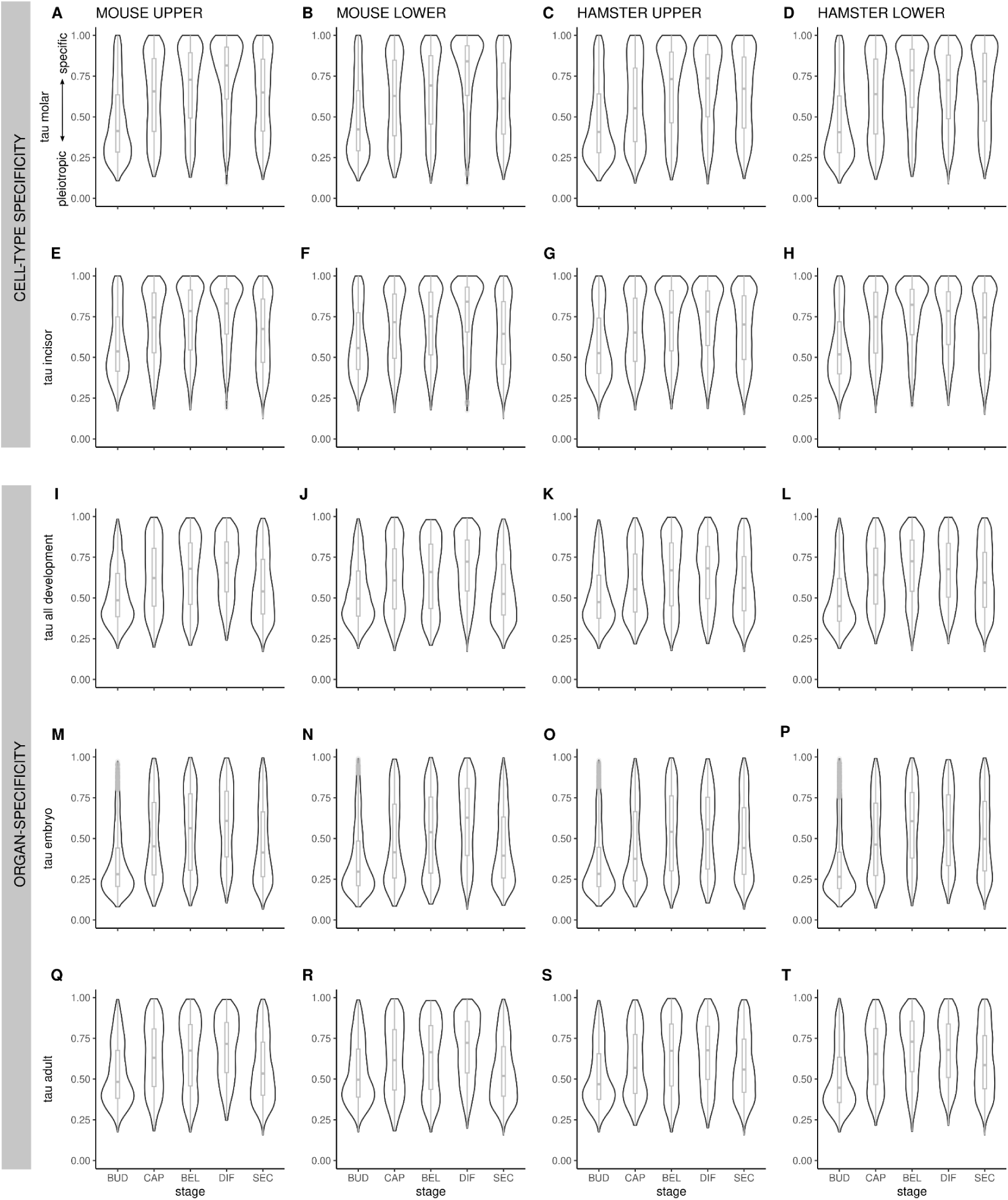
Distribution of expression specificity calculated by the tau index, in different data sets, for the 4 molars. Cell expression specificity was measured on single cell RNA-seq data in mouse molar (A-D) and mouse incisor (E-H). Organ/stage expression specificity was measured on the Bgee data for all development (I-L), embryonic stages (M-P), and adult stages (Q-T). It is represented by a violin plot for each stage cluster. Boxplots represent the median and the lower 25% and top 75% quantiles. Tau values range from 0 (pleiotropic) to 1 (organ/cell specific).

Finally, we also computed tissue-specificity based on a meta-analysis of a large number of organs present in the Bgee database [70]. We found similar results to those from the molar data, with genes in coexpression clusters peaking at early and late stages being more pleiotropic (here: expressed in many stages and organs) than genes peaking at mid stages (Fig. 7).

### Transcriptomic Age Index is lower at early stage, and hides composite contribution at late stage

We implemented a Transcriptomic Age Index (TAI [13]) that combines the age of genes with their level of expression. Expression levels were log transformed to obtain stable estimates and minimize impact of outlier values [19]. We observed lower values for the BUD stage than for the rest of morphogenesis (p<10-16, Fig 8C). We then separated the contribution of genes to TAI by phylostratum. We observed that the overwhelming majority of genes expressed in molars belong to the oldest strata, and are pleiotropic genes (in upper molar, 10716/14258 = 75% belong to PS1-PS2, and 87% are older than bilateria, Fig 8). Only more recent strata, corresponding to vertebrates and younger, carry a signal that varies across stages, which is in line with the evolutionary origin of the tooth. The transcriptome at the BUD stage proportionally lacks more recently evolved genes, which aligns with the presence of many conserved and pleiotropic genes involved in the development of ectodermal appendages.

**Fig 8.**
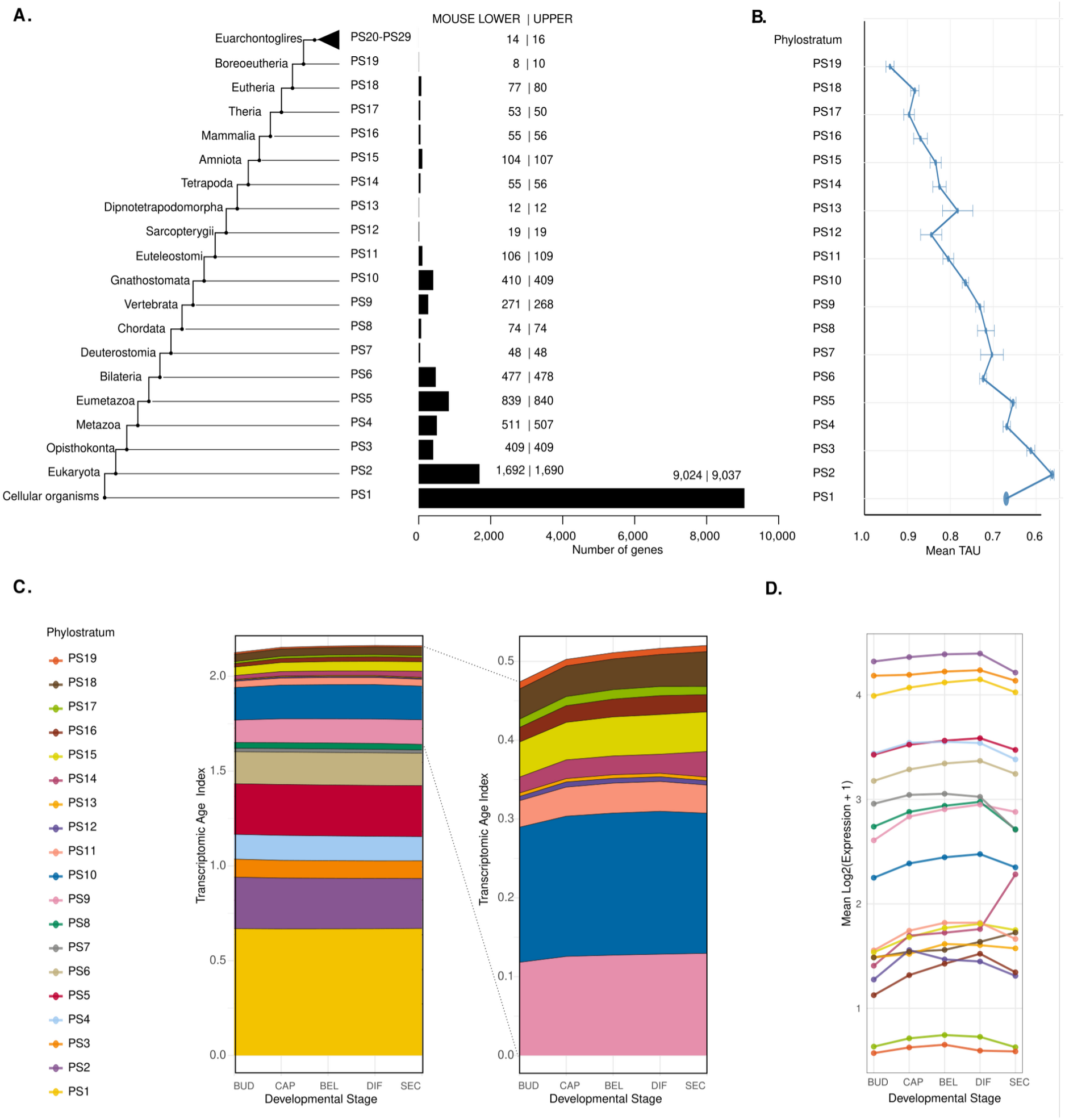
Age of genes expressed at different stages of molar development. (A) Number of genes expressed in lower and upper molar in mouse associated with each phylostratum. Younger phylostrata contained too few genes and were merged (above PS20). (B) Average pleiotropy measured for genes per phylostrata, up to PS19. (C) Left: Contribution of each phylostratum to the global TAI (Transcriptome Age Index). Right: Zoom for the contributions of phylostrata PS9 and younger. (D) Average expression for each phylostratum.

Even though the TAI value for the SEC stage was comparable to those for the earlier stages, we found that the age of transcriptome was more composite, with younger genes –particularly those from phylostrata 13 and younger, which correspond to tetrapodes– having a greater influence (Fig 8D). This could correspond to the onset of mineralisation genes, which are expressed at the SEC stage (Fig 2B) and whose evolution has been linked to modifications in bone and enamel associated with the transition from water to land.

## DISCUSSION

By applying different indexes to measure molar development conservation between mouse and hamster, we observed different patterns, which capture the complexity of the interplay of evolution and development. These patterns are composite and reflect the effect of both direct and indirect mechanisms that change during development. Also, we observed that the same pattern may be observed with different indexes and have different causes (Fig 9).

**Fig 9.**
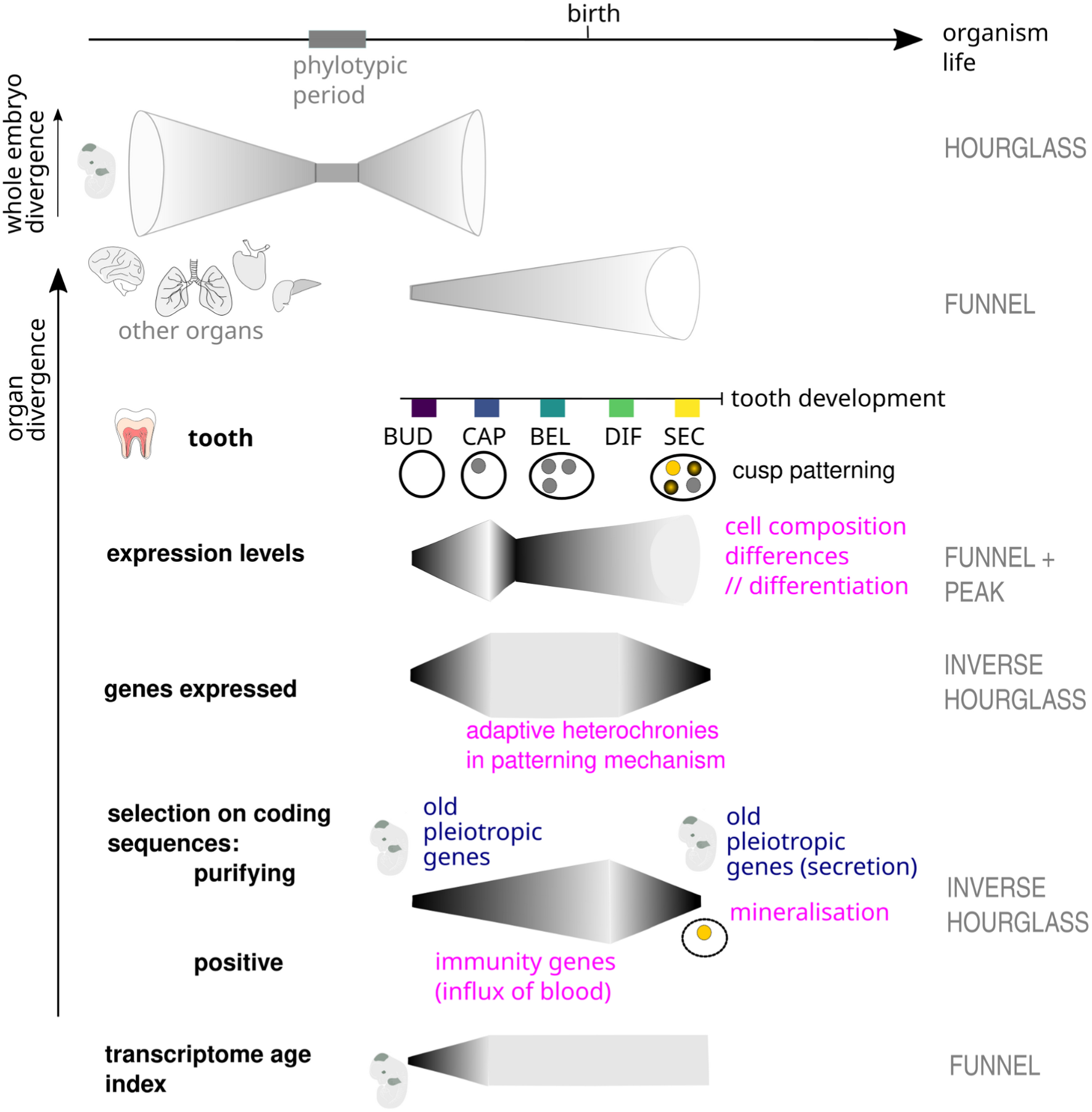
The conservation of the expression and sequence of genes expressed in molar development forms different patterns, attributable to processes shared with other organs or intrinsic to the tooth. Stages of tooth development are indicated, in parallel with the timing of patterns of conservation measured in other organs and in whole embryos. Shared (pleiotropic, blue) and intrinsic (tooth-specific, pink) processes that were linked to the patterns observed with different measurements in the manuscript are indicated. Development is not to scale.

Patterns found with the rate of coding sequence evolution form an inverse hourglass pattern which is consistent with assumptions based on expression and pleiotropy levels. The relationship between the rate of sequence evolution and the level of gene expression has been long discussed [71–76]. Highly expressed genes, genes with conserved temporal expression profiles, and genes that are pleiotropic—that is, expressed in a variety of cell types and organs—show on average the most intense purifying selection on coding sequences. These characteristics are often correlated with each other, for biological and technical reasons. By isolating these effects, we demonstrate that the coding sequence of genes exhibiting expression peaks at BUD and SEC stages display more purifying selection for different reasons. Genes whose expression peaks in early molar development (BUD) are pleiotropic since they are expressed in more tooth cell types and in more adult and developmental organs. Pleiotropy puts constraints on the evolution of coding sequences and has been linked before with periods of conservation of development [21,27]. By contrast, while some of the genes whose expression peaks in late molar development (SEC) are highly pleiotropic, others have high intra-cellular levels of expression, and are not as pleiotropic. Several models have been proposed to explain why highly expressed genes evolve slowly, including mistranslation, protein misfolding, and protein misinteractions [77–79]. This could apply to the SEC stage, when intense cellular matrix production, deposition and mineralisation occurs. We see that genes necessary for enamel, dentin and alveolar bone deposition start being massively expressed at this stage (S26 Fig). Because new genes more specifically involved in tooth and bone mineralisation evolved with water to land transition, but others involved in matrix production and secretion represent a very ancient toolkit, this may explain why the pleiotropy signal and the transcriptome age index are composite at SEC stage.

The amount of positive selection on coding sequences also creates an inverse hourglass pattern, with a peak at the DIF stage in mouse and a plateau in the BEL/DIF/SEC stage in hamster. Individual genes showing traces of positive selection at these stages are not particularly associated with known processes of molar development, which does not support a specific association with adaptive changes in tooth shape in rodents. On the contrary, they are enriched in immune-related genes, in particular the interferon pathway. The discovery of immune-related genes is common in analyses of positive selection, because these genes are at the forefront of host-pathogen interactions [80]. We believe that we find those genes in the late stages, because following the late cap stage, blood supply is gradually set up and immune cells are recruited locally in the pulp tissue [81]. Immune-related genes will therefore peak in late stages, and bring with them their positive selection signal. Although those immune cells and the control of inflammation in the pulp are surely important for tooth function [82], these cell types are also important elsewhere for organism immunity. In conclusion, the pattern of positive selection is largely driven by local recruitment of cells under selection for their function elsewhere. Depending when an organ is colonized by immune cells, its pattern of molecular evolution will be affected.

Expression divergence patterns are more challenging to understand than coding sequence divergence patterns because they differ based on whether one looks at the conservation of expression levels (which show a funnel with a local peak at CAP stage) or the conservation of expression peaks (which show an inverse hourglass). We think this is because expression divergence measured in bulk tissues incorporate variations in the nature and relative abundances of cell types, each having distinct effects on the transcriptome [83]. Divergence in the relative abundance of cell types only impacts the relative expression levels of the genes, but divergence in the nature of cell types affects both the sets of expressed genes and their expression levels. Overall conservation at BUD stage may therefore be interpreted as a conservation of cell types (sets of genes peaking at BUD stage) and their relative abundance (expression levels). At SEC stage, we observed that the set of genes was conserved but their expression levels were divergent. Hence, we postulate that the same cell types (in particular secreting cells) are present in both species, but their relative abundance differs. Because cusp patterning is more gradual in mouse molars than in hamster molars ([45]), we expect that a cell composition offset starts accumulating from BELL/DIFF stage, in terms of proportion of differentiating and then secreting cells, and culminate at SEC stage, when these processes are still ongoing.

We observed a stronger divergence of gene sets at mid stages of molar morphogenesis (CAP), that may carry hallmarks of adaptation. The divergence is even stronger for the upper than for the lower molar (Fig 3, S8 Fig). This coincides with an extended period of lingual growth specific to mouse upper molar, that was linked with the development of its new phenotype [45]. This relates to a period of divergence similarly placed during the intermediate stages of plant inflorescence development [84,85]. Interestingly, in both plant inflorescences and molars, mid-development involves the patterning of iterated structures in a growing bud, namely flowers in a growing meristem and cusps in a growing tooth bud (from CAP to DIFF stage for molars). In both cases, heterochronies in this process were linked to differences in final morphology [44,45,86,87]. This opens the possibility that minimal transcriptome conservation is observed at mid-development in such systems, because more heterochronies are present there, in relation to morphological adaptation, and that these blur the conservation signal in a process that would otherwise be largely homologous. In agreement with this hypothesis, we observed that genes peaking at mid-development tend to transition to the neighboring stage more frequently (Fig 3A-B; BEL <-> DIF transitions, with culmination of transcriptomic divergence at the DIFF stage).

Beyond the usage of same cell types at homologous stages, BUD stage may be particularly conserved because genes expressed there belong to processes particularly pleiotropic. Many processes involved in early tooth bud morphogenesis are shared with other skin appendages such as limbs, scales or glands, but also with lung or kidney branching morphogenesis [88,89]. Teeth and scales are ancestral to all jawed vertebrates and limbs are ancestral to tetrapods, and the components of the underlying developmental programs represent a very ancient molecular toolkit [90,91]. This program is even further recycled in other instances in body development, as demonstrated for mammary gland in mammals or patagium outgrowth in bats and sugar gliders [92]. Such ancient homology results in a high degree of pleiotropy, which is an important indirect source of evolutionary constraint, exposing these morphogenesis genes to stronger negative selection both on their expression in teeth and on their coding sequences.

To conclude, in all analyses, the BUD stage was most conserved with a progressive decline of developmental conservation from this early stage. When looking at the patterns of evolution for individual organs, sampling can only start when organs become available for dissection (e.g., around E10.5 in mouse), that is, after the onset of organogenesis, when the embryo is just coming out of the phylotypic period of the developmental hourglass model waist. Hence the evolutionary pattern observed for molar morphogenesis fits with the general interpretation that gene regulatory complexity is higher in mid-development, and late development is more permissive to the fixation of mutations. It also makes sense with respect to the nature and age of the processes involved. Due to the tooth bud simplicity and to this use of ancient genetic toolkits, everything converges for all indices to have low values at the BUD stage. Later, the signal is more composite and varies depending on the index employed, as the organ complexifies with additional cell types, differentiation, and the arrival of external cell types. The interpretation of diametrally opposed divergence observed for SEC stage (conserved in terms of gene set, but divergent in terms of gene expression levels) led us to highlight that this stage corresponds to the deployment of very conserved processes but with different magnitudes in different species. Finally, we also demonstrate that positive selection in coding sequences and sets of genes peaking at particular stages follow a similar inverse hourglass pattern but for very different reasons, some extrinsic to the tooth. By chance, colonization of immune cells in the tooth happens at the same time as heterochronies in cell type differentiation related to tooth morphological change. Before drawing the conclusion that an index only assesses the conservation of development in the organ in question, it is therefore important to analyze its meaning and exercise caution.

## METHODS

### Ethic Statement

This study was performed in strict accordance with the European guidelines 2010/63/UE and was approved by an Animal Experimentation Ethics Committee named « CECCAPP » (Lyon, France) and registered under # C2EA15 by the Ministry for education and research “Ministère de l’Enseignement Supérieur et de la Recherche”.

### Sample preparation

A total of 103 molar samples were obtained, corresponding to upper and lower molars from staged embryos in hamster and mouse. 36 samples were collected specifically for this study to complete our previously published data [47] (S1 Table). They cover, in duplicates, the whole period of tooth development in mouse from embryonic days (E) E12.5 to E22.5, equivalent to postnatal (PN) PN2 and in hamster (from E11 to E19.5, equivalent to PN2). Each sample contains two whole tooth germs, the left and right first molars (M1) of the same female individual (embryonic gonads were dissected to check for sex), and for a given stage, upper and lower samples were prepared from the same individual. Dissections were as tight as possible to include only M1 tissue, excluding the posteriormost molar prospective regions (as in [44]). The heads of harvested embryos were kept for a minimal amount of time in cooled advanced DMEM medium (small scale) or advanced DMEM medium (large scale). The M1 lower and upper germs were dissected under a stereomicroscope and stored in 200-500µl of RNA later (SIGMA), adjusting for sample size. Total RNA was prepared using the RNeasy micro kit from QIAGEN following lysis with a Precellys homogenizer. RNA integrity was controlled on a Bioanalyzer or Tapestation (Agilent Technologies, a RIN of 10 was reached for all samples). PolyA+ libraries were prepared with Illumina stranded mRNA prep kit), starting with 150 ng total RNA as in the previous study, and sequenced on an Illumina Hi-seq4000 sequencer at the Lausanne Genomic Technologies Facility. Because previous libraries had been generated with the non-stranded Illumina protocol and sequenced with a different design and platform, we also included 4 technical replicates, one per each end of the timeseries. We obtained on average 44 millions reads per sample with 150bp single-end strand-specific data for 40 samples (batch 2 and 2.1 for technical replicates, S1 Table) and 100 bp paired-end non-strand specific data for 65 samples (batch 1).

### Expression levels

These reads were mapped by using STAR (version 2.7.3a [93]) to reference sequences for golden hamster and house mouse transcriptomes. To generate them, we retrieved mouse and hamster genomes and annotations from Ensembl (release 98, January 2020, assemblies GRCm38 and MesAur1.0, [94]). The number of reads per genes was obtained by STAR - quantMode GeneCounts option, with default settings accounting for the strand-specificity of the data. The mapping rate is slightly higher in mouse than in hamster, with on average 80% of reads mapping unambiguously to annotated genes in mouse, and 65% in hamster. A table of counts and of TPM (transcripts per millions) values were created by a custom script based on these raw counts and on transcript length taken from Ensembl (release 98).

### Orthology relationships

Pairs of one to one orthologs between mouse and golden hamster were retrieved through Ensembl (release 98) by using biomaRt R library (version 2.48.0, [95]). Among these genes, only those with a MGI identifier were kept for further analysis. We kept 15910 pairs of orthologs.

### Multivariate analysis and global transcriptomic distances

The total table of raw counts contained 11342 genes with 1:1 orthologs in mouse and hamster, and with expression data in tooth development. We removed the samples corresponding to the upper and lower molars of one outlier individual (hamster E11.5), retaining 103 samples. We took the log of TPM counts and normalised these values with normalizeQuantile from the limma package [96]. We implemented PCA using the prcomp function from the stats package. We measured pairwise distance between pairs of samples of the same stage by using Spearman correlations.

To estimate the heterochronies, we used the fact that log transformed embryo weight is well correlated with its developmental age (especially when selecting only females, as we did here). By using a previously established linear relationship, we estimated development age from weight, and used samples aligned in a previous study [47,49] to align and scale time estimates between species (these samples represent times 0 and 10). We also obtained an estimate of developmental time derived from the second principal component of the PCA made on expression levels.

### Clustering of coexpressed genes

Stage-specific genes: Bgee Call version 1.4.0 was used with default parameters to generate present/absent gene expression calls [97]. Ensembl (release 84) was used for the mouse reference intergenic sequences. Treatment of transcriptome and genome data in the Bgee database include the definition of reference intergenic regions, which have very low probability of being false negatives of gene or exon annotation. For species in Bgee, we define reference intergenics as part of the database. For species not in Bgee, to apply BgeeCall, we need to define reference intergenics separately, and we store them in Zenodo (https://zenodo.org/communities/bgee_intergenic/).

Clusters of coexpressed genes: For each tooth type, the corresponding table of raw counts was first normalized with DESeq2 with the sequencing replicate as a batch effect (3.11, [98]). EISA clustering (eisa 1.44.0, [99]) was made with random starting clusters of samples (“random seeds”) and guided starting clusters of samples (“smart.seeds”). In the later case, starting conditions were obtained by grouping samples per developmental stage (bud, cap, bell, differentiation, secretion). The function ISAIterate was used to optimize the samples and genes grouping into clusters with the following parameters : thr.feat=1, thr.samp=1, convergence=“cor”. In the output, each gene has a score for each module. We assigned a gene to a particular module if the score of this gene is maximum in this module. Genes with a null score in all modules were not assigned.

### Functional enrichment

Functional enrichment was performed with the clusterProfiler [100] and ReactomePA [101] packages, using the enrichGO function to compute enrichment analyses in biological processes (BP) and molecular functions (MF) and the enrichPathway function to perform pathway enrichment analysis. To obtain a broad overview of the functional categories which are over-represented in gene sets, all functional enrichment analyses were performed with p.adjust < 0.1 and a maximum of 50 categories were reported. Terms were sorted by fold enrichment computed as following: FoldEnrichment = GeneRatio / BgRatio, with GeneRatio = k/n and BgRatio=M/N; with n genes in a given coexpression cluster, M genes in the functional gene set considered, k the overlap between the coexpression cluster and the functional gene set and N the number of unique genes in the dataset.

### Modelling gene expression profiles

Expression profiles were fitted by third degree polynomial splines with one interior knot, for each tooth separately species, using DESeq2 [98] and bs function of spline R package [102], as in [45]. To compute the distance between pairs of temporal expression profiles, we fitted a model with one specific curve per tooth. We computed for each gene the values predicted by each tooth model for 100 equally distributed points on the timeline. Using these profiles, we computed the time of the maximum expression level for each tooth. We also split them into 9 sliding time windows and measured the Euclidean distance point by point on each window. Then, for each gene we ranked the windows from the lowest to the highest distance.

### Rates of sequence evolution

Selective pressure on coding sequences, estimated by dN/dS between mouse and golden hamster, were retrieved for 14382 pairs of one-to-one orthologs through Ensembl (release 98) by using biomaRt R library (version 2.48.0).

Estimates of positive selection were extracted from the Selectome database (V7, based on Ensembl release 98, (Moretti et al. 2014; Proux et al. 2009)). For each gene, and for a selection of branches of interest, genes with a strong signal of positive selection (q-value < 0.2) were extracted and their functional enrichment was studied with StringDB [103]. We called these “genes with strong signal” despite the high q-value threshold because most genes have only marginal signal of positive selection, given the low power of the branch-site test.

We also obtained the likelihood ratios of models of H1 to Ho models with and without positive selection, from the branch-site model [104]. The obtained ratios, Δln*L,* represent the evidence for positive selection. We transformed Δln*L* with the fourth root to improve their normality, following [58,59,105]. We obtained data for 11,460 genes on the Muridae branch, 12,919 genes on the murinae branch, 13,257 genes on the *Mus* branch, and 10,936 genes on the Cricetinae branch.

We split genes according to their phase of expression, following the clusters defined by EISA. Then, we computed the mean of dN, the mean of dS, and took their ratio mean(dN)/mean(dS). We also directly took the mean and median dN/dS of the genes per cluster. To get an estimate of the uncertainty of these values, we bootstrapped gene contents by resampling genes with replacement within clusters 1000 times, and recomputed the mean/median dN/dS as above. The same procedure was used for the 4th root of likelihood ratios taken from selectome.

To include all genes expressed at each developmental stage, and inspired by [27] we computed an index of purifying selection on coding sequence as 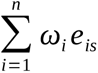

where s is the developmental stage, *ω*_i_ is the dN/dS of gene i, n is the total number of genes, and e_is_ is the expression level of gene i in developmental stage s; we use log-transformed expression levels for e_is_.

For obtaining confidence intervals for these data, we represented error bars, and as an alternative, we performed bootstrapping (still following [27]). For this, we randomly sampled gene IDs from the original data set 10,000 times with replacement. Then, we computed transcriptome indexes for the 10,000 samples. Finally, we defined 95% confidence intervals as the range from quantile 2.5% to quantile 97.5% of the 10,000 transcriptome indexes.

### Immune-related genes

We extracted a list of 3094 mouse genes that belong to the gene ontology term “immune system process” (GO:0002376, https://www.informatics.jax.org/vocab/gene_ontology/GO:0002376). 1512 of these genes are part of our dataset because they have orthologs in the hamster. To check the consistency of our results, we removed them and redid the analyses on the remaining 9830 genes.

### Pleiotropy/Tissue specificity

To estimate pleiotropy/tissue-specificity at a multi-organ and multi-stage scale, we extracted expression values for three datasets: all developmental stages, embryonic and post-embryonic mouse libraries from the Bgee database (version 15.2) in September 2024 [106]. We log transformed the expression values (TPM) and calculated the mean of the log transformed expression values per anatomical feature. We obtained data for 117 organs and 55,486 genes (all developmental stages); 38 organs and 55,410 genes (embryonic stage), 86 organs and 55,486 genes (post-embryonic).

To estimate pleiotropy in teeth, we extracted expression values from two published single-cell RNA-seq datasets from wild type and healthy mice, one in incisors and one in molars. In incisors, we obtained expression levels for 2889 cells and their annotations in 17 cell types (GSE146123, [69]. In molar, we obtained 30930 cells from E14.5 stage lower molars, annotated in 23 cell types (GSE142200, [68]). We used Seurat package to obtain pseudo bulk expression profiling for each cell type [107].

We estimated pleiotropy in these 5 datasets by calculating Tau as previously described [108]. This index takes into account the mean of the log transformed expression values and the number of organs considered. Tau was calculated as: τ = sum(1-xb)/(n-1) where n is the number of organs considered, and xb=x/xmax is the level of expression normalized by the maximum level of expression in the vector. τ ranges from 0 (ubiquitously expressed) to 1 (expressed in a single organ/developmental stage).

### Phylostratum temporal expression profile analysis

Mouse and hamster gene expression values for mandible and maxilla tissues were log transformed as log₂(TPM + 1) and averaged across replicates per gene for each of the 5 developmental stages, BUD, CAP, BEL, DIF, and SEC. Genes were converted to protein IDs then mapped to the Mus musculus phylostratum reference using the package phylomapr (v. 0.0.2, using https://github.com/LotharukpongJS/phylomapr and the GenEra data [109]). For hamster, we first mapped hamster genes to mouse orthologs followed by assignment of evolutionary age.

We analysed the expression pattern of each phylostratum across development stages by computing the mean expression of evolutionary gene sets at each development stage only including phylostrata with a minimum of 5 genes. With this approach we decomposed the aggregation of different patterns to reveal the contribution of each gene set at different developmental stages. The analysis was carried out separately for upper and lower molar per species.

### Phylostratum-based pleiotropy analysis

We obtained mouse gene expression data across all developmental stages and 24 tissue types from the Bgee database (version 15.2) using the R package BgeeDB (v. 2.31.2). We separately analysed all developmental stages, embryonic (UBERON:0000068 and descendants), and post-embryonic stages (UBERON:0000092, UBERON:0000113 and descendants). We first log transformed the TPM expression values as log₂(TPM + 1) and averaged across replicates for every gene-tissue combination. Each gene was assigned an evolutionary age by converting Ensembl gene IDs to protein IDs using biomaRt (v. 2.58.2) then mapping to the Mus musculus phylostratum reference (Mus_musculus_ENSEMBL_GRC38mm10.PhyloMap) using the package phylomapr (v. 0.0.2). We then calculated the Tau index for each gene across all tissues as previously described.

To test the hypothesis that older genes are more broadly expressed than younger genes, mean Tau values and standard errors for mean Tau within each phylostratum were computed for each phylostratum with only phylostrata with a minimum of 5 genes being included in the analysis.

## Data availability

Raw data are publically available in ENA with project accession numbers PRJEB52633 and PRJEB84925. Data is available in Zenodo, DOI 10.5281/zenodo.15374260.

## Code availability

All custom code (run in R) used in this study is made available here: https://gitbio.ens-lyon.fr/LBMC/cigogne/molar_inverse_hourglass

## ACKNOWLEDGEMENTS

We acknowledge the contribution of several platforms of SFR Biosciences Gerland-Lyon Sud (UMS344/US8): the Plateau de Biologie Expérimentale de la Souris (PBES) (many thanks especially to Jean-Louis Thoumas, Tiphaine Dorel, Céline Angleraux, Marie Teixeira, Myriam Prudent), as well as the computer resources from CBPSMN (ENS Lyon).

We acknowledge the technical help of Anne Lambert, Alain Rubod, Mathilde Estevez-Villar, and the contribution of many students including Coraline Petit, Alice Lorenc, Margaux Pillon, Ludivine Rotard and Asma Benahmed.

## AUTHOR CONTRIBUTIONS

Data curation : JG, SM; Formal analysis : JG, SM, MN, MS; Investigation : JG, MEV, MM, SM, MN, SP, MS; Methodology : JG, MRR, SP, MS; Visualization : JG; Writing – review & editing : JG, MRR, SP, MS; Writing – original draft : SP, MS; Project administration : MRR, SP, MS; Conceptualization : MRR, SP, MS; Funding acquisition : MRR, SP, MS; Supervision : SP, MRR, MS.

## LIST OF LEGENDS TO SUPPORTING INFORMATION FILES

S1 Data. Matrix of raw counts

S2 Data. Matrix of TPM values

S1 Table. Description of the samples used in the study.

S2 Table. Number of stage-specific genes for each tooth and species.

S3 Table. Comparison of dN/dS values between genes with a conserved and non-conserved expression peak, per stage and for different comparisons.

S4 Table. Genes with evidence of positive selection. Stage indicates corresponding coexpression cluster (taken from hamster upper molar for the cricetinae branch, and from mouse upper molar otherwise). q values are indicated.

S1 Text. Sampling and heterochronies.

S1 Fig. Comparisons of relative times of development for both species. (A) Relative times estimated from weight and in days post coïtum are coherent between species, with distinct periods for each morphological stage (shaded bands). (B) development time estimated from embryonic weight and from the second axis of a PCA analysis. Some samples fall into the range of developmental age normally occupied by the previous or next morphological stage (highlighted). This concerns 4 samples in hamster (2 BUD, 1 CAP, 1 DIF). Morphological stages are colored, species and tooth types are indicated with different shapes.

S2 Fig. Main effects in the tooth developmental transcriptome. (A) PCA based on 11,342 1:1 orthologs depicted in Fig 1A (each dot represents a single tooth) with embryonic days indicated (dpc), samples colored per morphological stage, shapes corresponding to upper and lower molar samples. (B) PC1 and 3 of the same PCA with symbols representing the batch of sequencing. (C) scree plot describing the amount of variance explained by the first 10 PCs.

S3 Fig. Pictures of the lower molar samples used for preparing the RNAseq secretory stage in our dataset. Stage (days post birth and days post coïtum) and embryo weight (in mg, all embryos are female) are shown. Mineralisation progression is visible in each species, starting from the tip of the cusps down into the valleys (white arrows). Mineralisation at PN0 in hamster might be slightly closer from mineralisation at PN2 than PN0 in mouse. However, physiological changes at birth may also impact the transcriptome, so we considered taking PN0 and PN2 in both species was a good compromise.

S4 Fig. Summary of the functional enrichment analysis for the five coexpression clusters in lower mouse molar; enriched Biological Processes GO terms were selected for their adjusted p-value (<0.1) and sorted per fold enrichment (cluster gene ratio/background ratio, see Methods). A maximum of 50 GO terms are represented.

S5 Fig. Summary of the functional enrichment analysis for the five coexpression clusters in lower mouse molar; enriched Reactome Pathways were selected for their adjusted p-value (<0.1) and sorted per fold enrichment (cluster gene ratio/background ratio, see Methods). A maximum of 50 GO terms are represented.

S6 Fig. Overlap of coexpressed gene clusters between molars of the same species. Each cluster corresponds to a set of genes with an expression peak at a specific stage of development in a given tooth and species. Number of genes shared between hamster and mouse coexpression clusters, expressed as a fold enrichment relative to equally sized random clusters, for mouse (A) and hamster (B). Clusters are ordered according to their temporal profiles. Proportion of genes shared between upper and lower coexpression clusters peaking at the same stage in both teeth, in the mouse (C) and hamster (D) molars. Intervals show the 95 percentiles obtained by 1000 bootstrap resamplings. Expected values obtained by randomly shuffling genes between clusters (dashed lines)

S7 Fig. Curves modelling gene expression allow to account for batch effect (A), extract time of maximum gene expression for each tooth profile (B) and measure local divergence in 9 sliding windows and rank them (C). Example is shown for the Shh gene in the upper molar.

S8 Fig. Offset between peaks of maximal expression. Gene temporal expression profiles were modelled individually by splines with one curve per species. The peaks of expression were computed for each species. Related to Fig 3A-D

S9 Fig. Local divergence of expression levels peaks at CAP and SEC stages. Gene temporal expression profiles were modelled individually by splines with one curve per species. The area between curves was computed in 10 sliding windows along development. These windows were ordered from the most conserved to the most divergent for each gene. For each stage, we then obtain a measurement of the relative conservation of expression levels. Related to Fig 3E.

S10 Fig. Distribution of dN/dS ratio, over 14382 mouse genes with one to one orthologs in hamster. The distribution was cropped to dN/dS=1, removing 22 genes. The mean of the distribution is indicated on the graph.

S11 Fig. Index of purifying selection on coding sequence calculated as the dN/dS of each gene weighted by its expression at each morphological stage. 95% confidence intervals around means are represented.

S12 Fig. Index of purifying selection on coding sequence calculated as the dN/dS of each gene weighted by its expression at each morphological stage. The result of 10000 bootstraps is represented, with means and quantile 2.5% to quantile 97.5%.

S13 Fig. Selective pressure acting on coding sequences in the gene coexpression cluster associated with developmental stages. Each panel represents one pairwise comparison between molars. Genes associated with the same stage in both molars are called “conserved” (green). Others fall in the “non-conserved” bin (red). In blue, total values (as in Fig 4). Each plot shows the average dN/dS computed as mean(dN)/mean(dS) over each gene set. a) Gene clusters in mouse upper molar compared with mouse lower molar. b) Gene clusters in hamster upper molar compared with hamster lower molar. c) Gene clusters in mouse upper molar compared with hamster upper molar. d) Gene clusters in hamster lower molar compared with mus lower molar. Observed values are represented by a line, intervals show the 95 percentiles obtained by 1000 bootstrap resamplings.

S14 Fig. Function of genes with strong evidence of positive selection (all branches pooled) analyzed with gene ontology biological processes (A) and with gene networks for early (bud/cap, B), mid (bell/differentiation, C) and late (secretion, D) stages. Number of genes is indicated. Networks were created with StringDB, only mid-development shows a significant network (links between genes) including the pro-inflammatory gene Interferon Gamma (Infg).

S15 Fig. Number of genes with sign of positive selection (q value <0.2) for each module and branch. The number of genes related to the immune system are highlighted in black.

S16 Fig. Index of positive selection on coding sequence calculated as the fourth root of the likelihood ratio (ΔlnL) of the models with and without positive selection, taken from Selectome database. The value for each gene was weighted by its expression at each morphological stage. The result of 10000 bootstraps is represented, with means and quantile 2.5% to quantile 97.5%.

S17 Fig. Positive selection for genes with expression peaks at different stages of development, in four branches of the rodent tree, after removal of immune-related genes. Each plot shows the fourth root of the likelihood ratio (ΔlnL) of the models with and without positive selection, taken from Selectome database. Average for coexpressed gene clusters associated with each stage of development are represented. Confidence intervals obtained from 1000 bootstraps. Branches used to compute positive selection are indicated on the rodent tree in A, and correspond respectively to B: Muridae, C: Murinae, D: Mus, E: Cricetinae. Gene clusters correspond to mouse upper molar (B-D) or to hamster upper molar (E). Species tree in panel A pruned from Ensembl Compara.

S18 Fig. Main effects in the tooth developmental transcriptome, without immune-related genes. PCA based on 9830 1:1 orthologs. Each dot represents a single tooth with embryonic days indicated (dpc), samples colored per morphological stage, shapes corresponding to batches of sequencing. PC1 and PC2 (A) and PC3 and PC4 (B) are represented. (C) scree plot describing the amount of variance explained by the first 10 PCs.

S19 Fig. Overlap of expression gene clusters between species, without immune-related genes. Each cluster corresponds to a set of genes with an expression peak at a specific stage of development in a given tooth and species. Number of genes shared between hamster and mouse coexpression clusters, expressed as a fold enrichment relative to equally sized random clusters, for the upper (A) and the lower (B) molars. Clusters are ordered according to their temporal profiles. Proportion of genes shared between hamster and mouse coexpression clusters peaking at the same stage in both species, in the upper (C) and lower (D) molars. Intervals show the 95 percentiles obtained by 1000 bootstrap resamplings. Expected values obtained by randomly shuffling genes between clusters (dashed lines).

S20 Fig Index of purifying selection on coding sequence calculated as the dN/dS of each gene weighted by its expression at each morphological stage. Immune-related genes were removed. 95% confidence intervals around means are represented.

S21 Fig Selective pressure acting on gene coding sequences for each coexpressed gene cluster, without immune-related genes. Average pairwise dN/dS between mouse and hamster were computed as mean(dN)/mean(dS). Observed values are represented by a black line, intervals show the 95 percentiles obtained by 1000 bootstrap resamplings.

S22 Fig. Correlation between dN/dS and expression level per stage. Genes were split by developmental stage according to their expression profile as in Fig 1. Expression was averaged for each gene, over the samples associated with their corresponding stage. Spearman correlations were computed between this average expression level per stage (in log(TPM)) and the dN/dS, for each cluster of genes. Observed values are represented by a black line, intervals show the 95 percentiles obtained by 1000 bootstrap resamplings.

S23 Fig. Selective pressure acting on coding sequences across developmental stages. Each panel shows, in black, the average dN/dS computed as mean(dN)/mean(dS) for groups of genes associated with different stages. In colors, mean(dN)/mean(dS) were computed separately on subsets of genes split by their level of expression. Observed values are represented by a solid line, intervals show the 95 percentiles obtained by 1000 bootstrap resamplings.

S24 Fig. Overlap of expression gene clusters between species, for quantities of low, medium-low, medium-high and high expression levels. Proportion of genes shared between hamster and mouse coexpression clusters peaking at the same stage in both species, in the upper (A) lower (B), mouse (C) and hamster (D) molars.

S25 Fig. relationship between the level of pleiotropy and the level of expression. Contour Plots represent the density of the distributions, for each molar and coexpressed gene clusters. Pleiotropy was computed with molar single-cell RNA-seq data.

S26 Fig. Expression levels for the Enamel gene obtained for the upper molars and color coded by morphological stage. Curves modelling gene expression allow to account for batch effect and show a sharp increase at the secretion stage.

